# Distinct Alpha Networks Modulate Different Aspects of Perceptual Decision-Making

**DOI:** 10.1101/2025.03.14.643170

**Authors:** Ying Joey Zhou, Mats W.J. van Es, Saskia Haegens

**Affiliations:** School of Psychology, Shenzhen University; Oxford Centre for Human Brain Activity, Department of Psychiatry, University of Oxford; Department of Psychiatry, College of Physicians and Surgeons, Columbia University; Division of Systems Neuroscience, New York State Psychiatric Institute

## Abstract

Why do we sometimes perceive a faint stimulus but miss it at other times? One explanation is that conscious perception fluctuates with the brain’s internal state, influencing how external stimuli are processed. Ongoing brain oscillations in the alpha band (8–13 Hz), thought to reflect neuronal excitability levels^1–5^ and play a role in functional inhibition^6,7^, have been shown as a key contributor to such perceptual variability^8,9^. Under high alpha conditions, faint stimuli are more likely to be missed^8^. Some studies suggested alpha oscillations modulate perceptual criterion (*c*)^10–14^, shifting the threshold for interpreting sensory information; while others (including our prior work^15^) suggested alpha modulates sensitivity (*d’*)^15–19^, changing the precision of sensory encoding. Few studies observed modulations in both metrics, making these results appear mutually exclusive. Most studies have focused solely on overall alpha activity— whether within a region of interest or across the whole brain—and overlooked the coexistence of multiple distinct alpha networks^20–26^, which fluctuate in terms of predominance^20,27,28^ and adapt to behavioural demands^29,30^. Hence, it remained unclear whether different networks’ contributions to perception vary with their momentary state. Here, aiming to characterize how different alpha networks influence perceptual decision-making, we analyzed magnetoencephalography (MEG) data recorded while participants performed a visual detection task with threshold-level stimuli. We found that while the visual alpha network modulates perceptual sensitivity, the sensorimotor alpha network modulates criterion in perceptual decision-making. These findings reconcile previous conflicting results and highlight the functional diversity of alpha networks in shaping perception.

## RESULTS

We analyzed MEG data recorded from 32 healthy participants while they performed a visual detection task involving ambiguous gratings (Figure 1A). Notably, we experimentally manipulated participants’ decision criterion via “priming” (Figure 1B). This manipulation allowed us to test whether top-down changes of criterion were implemented via (or reflected in) shifts in alpha dynamics. In our original work^15^, we operationalized alpha dynamics as the power and phase of alpha oscillatory activity in stimulus-responsive areas. In the current work, in contrast, we took a network perspective and aimed to link oscillatory alpha power in different large-scale networks to perception. We first applied the time delay embedded hidden Markov model (TDE-HMM)^20^ to the MEG source-level data to find large-scale brain networks in a data-driven way, in which we modeled the observed data as generated by eight recurring, transient, and mutually exclusive hidden states. Here, a state refers to a unique pattern of network activity, characterized by specific spatial and spectral profiles, across all the analyzed brain regions. Next, we identified states with prominent alpha oscillatory activity and defined the networks dominating these states as the alpha networks of interest. With these alpha networks defined, we then asked whether ongoing alpha power fluctuations in these networks modulate perceptual decision-making, and, if so, what the underlying mechanisms are.

**Figure 1.**
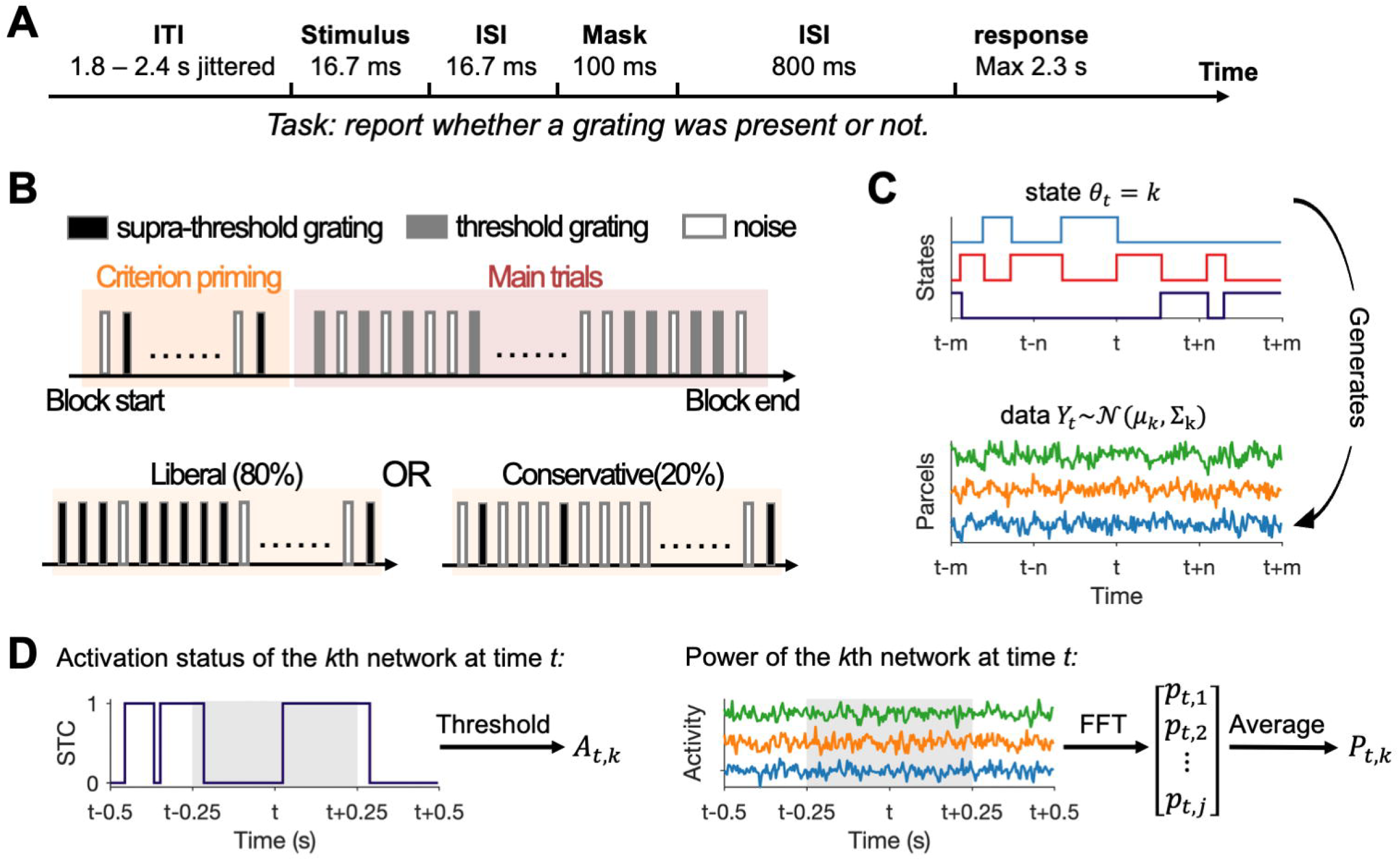
Experimental protocol and HMM analysis schematics. **(A)** Trial schematic of the main task trials. **(B)** Experimental procedure. **(C)** Generative model of the HMM. At each time point *t*, one state is active (e.g., the *k*th state), and the observed parcel-specific activity (data *Y*_*t*_) is assumed to be drawn from a state-specific multi-dimensional Gaussian distribution with mean _*k*_ and co-variance matrix _*k*_. **(D)** Schematics on how, for the *k*th network, its activation status at time *t* (*A*_*t*,*k*_) and network-level alpha power at time *t* (*P*_*t*,*k*_) were estimated. STC refers to the state time course. Grey shaded areas denote the 500-ms sliding window used.

### HMM reveals large-scale networks with distinct dynamics

The time-delay embedded Hidden Markov modeling returns temporally and spectrally resolved large-scale networks (Figure 2), each corresponding to a unique pattern of network activity (i.e., state). At each time point, one state is “on”, indicating which large-scale network dominates. The identified networks showed topography and connectivity profiles comparable with previous work^20,30^, even though the parcellation of source-level data in the current study was different. For example, state 8 exhibited positive wideband power and coherence in motor areas and negative wideband power in a wide range of areas covering the parietal-occipital part of the brain, consistent with the sensorimotor state reported by Vidaurre et al. (2018). At time points around stimulus presentation and around response probe onset, state 8 was significantly less likely to be active than during the baseline (*p* < 0.029 for the earlier cluster, and *p* < 0.001 for the late cluster; Figure S1A), further supporting its sensorimotor nature. In contrast, state 7 and state 4 exhibited similar wideband power and coherence maps to the “visual” and “posterior higher-order cognitive” states identified by Vidaurre et al. (2018). Moreover, state 4 and 7 were significantly more likely to be active in response to the stimulus relative to the baseline (*p* = 0.005 for state 4; *p* < 0.001 for state 7; Figure S1A). For simplicity, we will refer to states 4, 7, and 8 as the posterior higher-order cognitive state, the visual state, and the sensorimotor state, respectively; and the respective dominant networks in these states the posterior higher-order cognitive network, the visual network, and the sensorimotor network.

**Figure 2.**
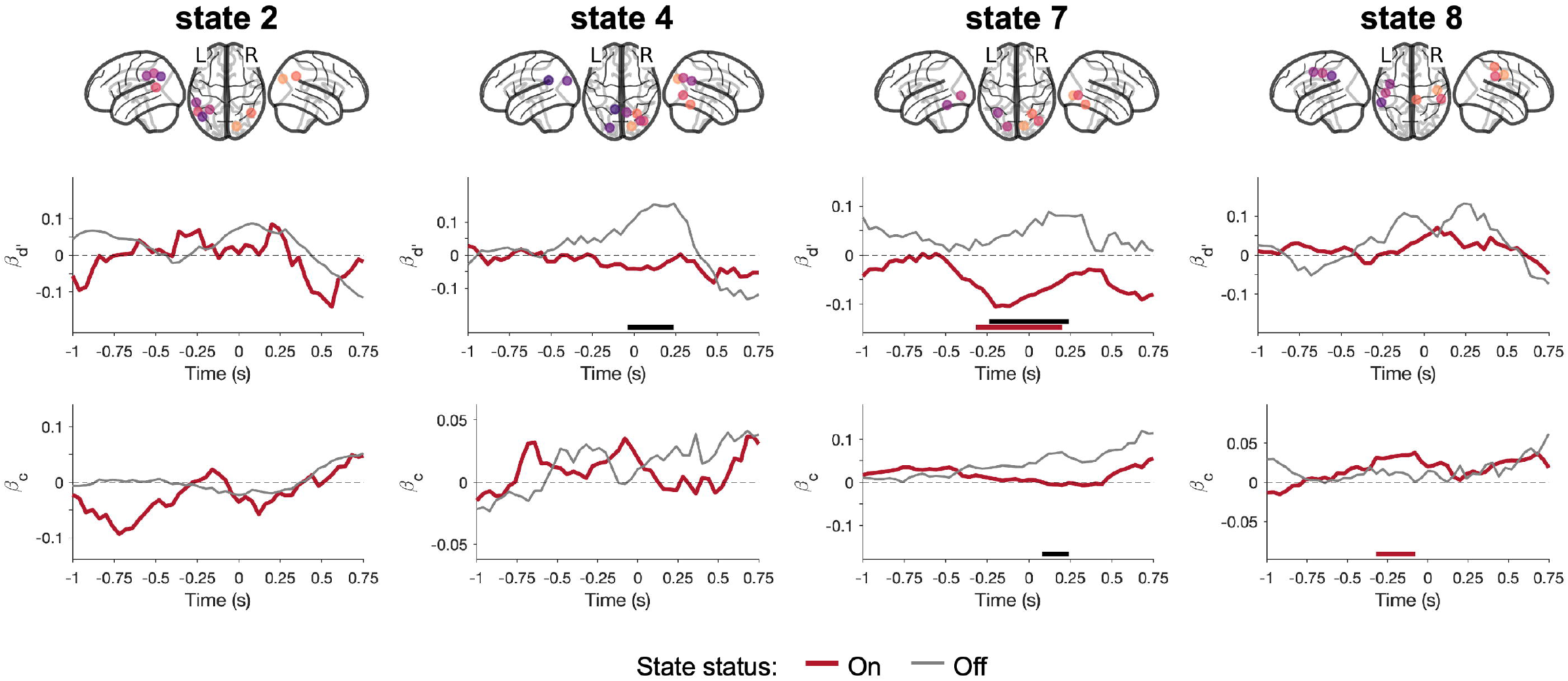
Networks identified by HMM. For each state, the spatial distribution of power and coherence were estimated for frequencies between 1 and 45 Hz. In each panel, the power map (top left) shows group-averaged power, relative to the mean across states. The coherence network (bottom) shows the top 98% coherence with colored lines, and centers of each parcel with black dots. The PSD graph (top right) shows both the state-specific (colored solid line) and static PSD (i.e., the average across states, black line).

### Spectral profiles distinguish different alpha networks

Next, we inspected the alpha-band power and coherence maps of the different states. We applied the FOOOF algorithm to the resulting group-average state-specific PSDs (i.e., PSDs computed over time points when the state is “on” and across the “critical nodes”). The algorithm returned peaks centered between 7 to 14 Hz for states 2, 4, 7, and 8, but not for states 1, 3, 5, and 6, suggesting that only states 2, 4, 7, and 8 exhibited a clear alpha-band oscillatory component (R^2^ > 0.99 for all models fit to the state-specific PSDs). We therefore focused on states 2, 4, 7, and 8 in the following analyses (Figure 3).

**Figure 3.**
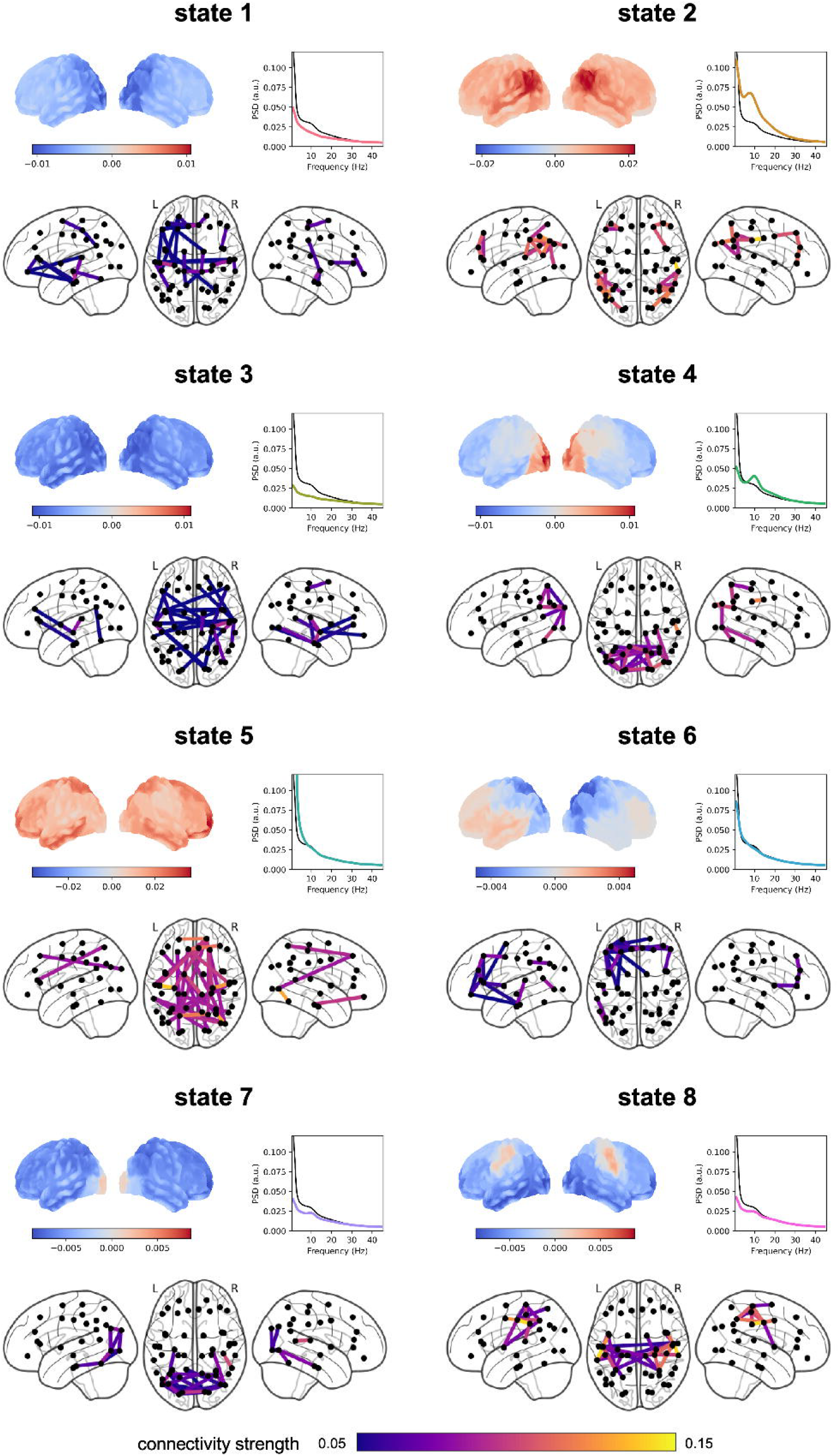
Networks exhibiting oscillatory alpha activity. **(A)** For each state, the spatial distribution of power and coherence were estimated for frequencies in the alpha range. Sub-panels are organized similarly to Figure 2; the PSD graph (top right) shows the group-averaged state-specific PSD in thick solid line (and participant-specific PSDs in thin lines) estimated from the critical nodes (highlighted in black in the coherence network). **(B)** The bandwidth and center frequency for each state as determined by the FOOOF algorithm. Dots denote individual participants (only those exhibiting an alpha oscillatory component were plotted).

To further establish the unique sources of alpha activity in these states, we asked whether the bandwidth and peak frequency of the alpha oscillatory components differ between these four states of interest. We applied the same FOOOF algorithm to the state-specific PSDs computed for each participant, obtained the fitted center frequencies and bandwidths within the 7 to 14 Hz range, and used repeated-measures ANOVA to test whether they differed across states. We excluded participants from the rmANOVA analysis if their alpha oscillatory component was absent in one or more states, leaving a total of 22 participants (i.e., 22 participants showed clear oscillatory alpha components in all four states of interest). Our results (Figure 3B) showed that the main effect of states on bandwidth was not statistically significant (*F*(3, 63) = 2.004, *p* = 0.122, Greenhouse-Geisser corrected *p* = 0.142), while the main effect of states on center frequency was significant (*F*(3, 63) = 6.541, *p* < 0.001, Greenhouse-Geisser corrected *p* = 0.003). Pair-wise comparisons on peak frequency showed that state 7 exhibited a significantly higher peak frequency (group-averaged center frequency *CF*_*7*_ = 11.02 Hz, alpha oscillatory component identified in *N*_*7*_ = 28 participants) compared to state 2 (*CF*_*2*_ = 9.83 Hz, *N*_*2*_ = 28; state 7 vs. state 2: *t*(26) = 3.659, *p* = 0.001), state 4 (*CF*_*4*_ = 10.31 Hz, *N*_*4*_ = 28; state 7 vs. state 4: *t*(27) = 3.447, *p* = 0.002), and state 8 (*CF*_*8*_ = 10.17 Hz, *N*_*8*_ = 25; state 7 vs. state 8: *t*(22) = 2.399, *p* = 0.025). Taken together, we identified four alpha networks of distinct spatial and spectral profiles, corresponding to four states.

### Ongoing alpha power fluctuation predicts different aspects of perception

Having identified the states and corresponding networks of interest (i.e., states 2, 4, 7, and 8), we then asked whether fluctuations of ongoing alpha activity modulate perceptual sensitivity. We addressed this question using a GLMM approach (see STAR Methods for details). In short, we modelled the participant’s behavioral report with the stimulus condition (grating present/absent), the network activity (on/off), the network alpha power, and their cross-terms. The obtained beta coefficients could be combined to reflect the extent of modulation of criterion (*β*_*c*,*A*=0_and *β*_*c*,*A*=1_) and sensitivity (*β*_*d*′,*A*=0_and *β*_*d*′,*A*=1_) over time (Figure 4). Concretely, a positive (or negative) *β*_*d*′_ at time *t* suggests that increased alpha power at time *t* leads to higher (or lower) perceptual sensitivity; a positive (or negative) *β*_*c*_ at time *t* suggests that increased alpha power at time *t* leads to more liberal (or conservative) criterion in reporting “target present”. Statistical significance of the beta coefficient time courses (one time course per state) was inferred using cluster-based permutation tests on the time window starting 500 ms before and ending 250 ms after stimulus onset.

**Figure 4.**
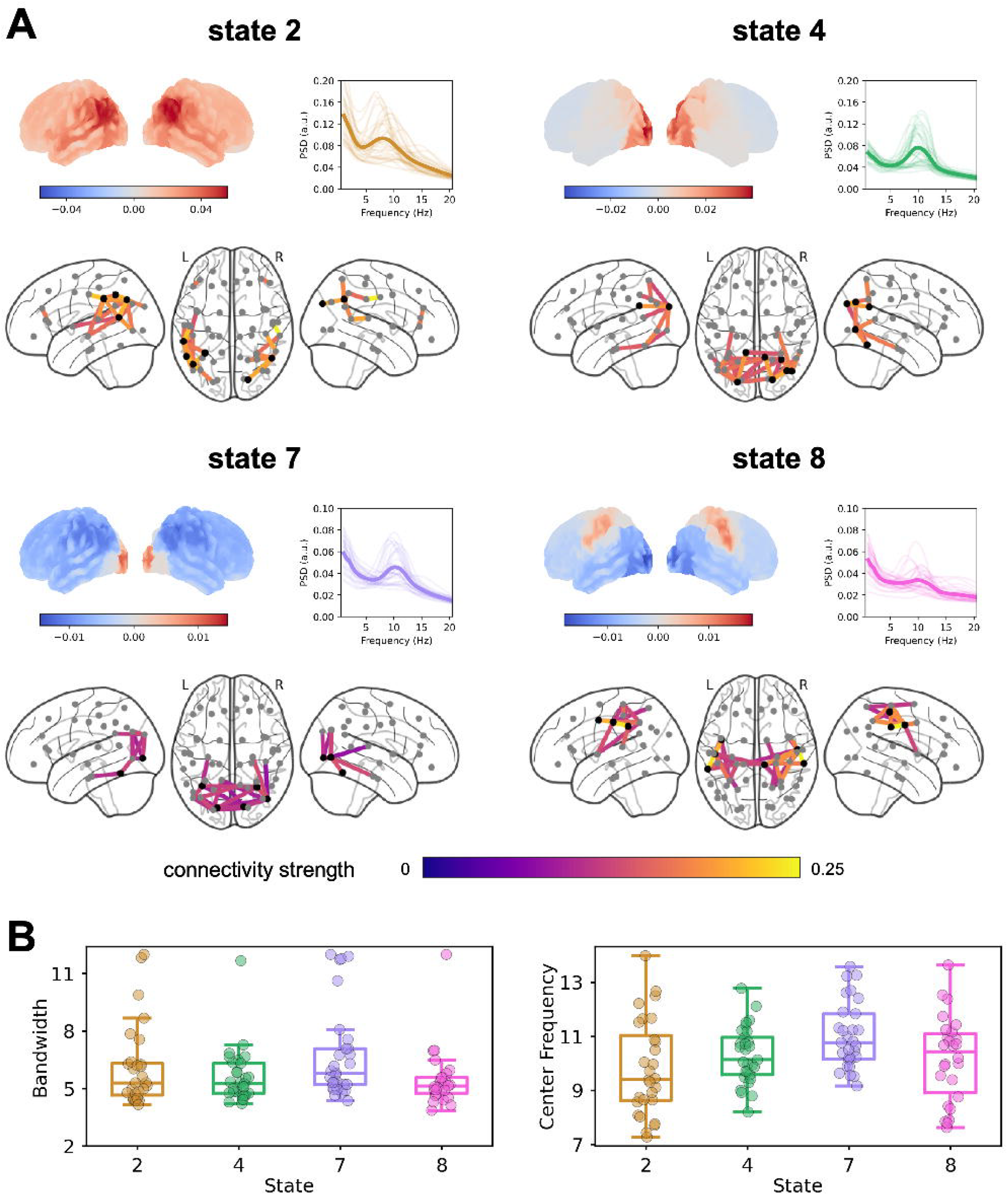
Linking alpha power to sensitivity and criterion. Top: The critical nodes used to estimate network-level alpha activity (highlighted in different colors for visibility). Bottom: Time courses of *β*_*d*′_ and *β*_*c*_ when the network was on (red) vs. off (grey). Horizontal bars indicate time points that contributed to significant effects (red denotes state activity (i.e., on/off), black denotes the interaction term (i.e., state on/off × alpha power)).

We first asked whether and how fluctuations of ongoing alpha activity modulate perceptual sensitivity (Figure 4). Our results showed that ongoing alpha power fluctuation in the visual network (i.e., the dominant network in state 7) significantly modulated sensitivity only when the state was on (*p* < 10^-4^), but not when the state was off (interaction effect *p* < 10^-4^). Crucially, the observed effects corresponded to clusters centering at time zero and expanding ∼200 ms both pre- and post-stimulus onset, suggesting that the sensitivity effect likely started pre-stimulus and lasted until after the stimulus presentation. Moreover, alpha activity fluctuation in the posterior higher-order cognitive network (i.e., the dominant network in state 4) displayed statistically different *d’* modulations when the state was on vs. when it was off (interaction effect *p* = 0.0006). This effect corresponded to a cluster covering mostly post-stimulus time points, suggesting that the interaction effect was mostly post-stimulus, likely caused by distinct stimulus-related activity when the state was on vs. off.

Similarly, we asked whether and how fluctuations of ongoing alpha activity modulate perceptual criterion (Figure 4). Our results showed that ongoing alpha power fluctuation in the sensorimotor network (i.e., the dominant network in state 8) significantly modulated criterion, especially when the state was on (*p* = 0.0458). This effect corresponded to a cluster around - 250 ms (locked to stimulus onset), suggesting it was most likely pre-stimulus. Our results also showed that alpha activity fluctuation in the visual network (i.e., the dominant network in state 7) modulated criterion differently when the state was on vs. when it was off (interaction effect *p* = 0.0124). This effect corresponded to a cluster around 150 ms, suggesting that this interaction effect was post-stimulus, most likely caused by distinct stimulus-related activity when the state was on vs. off.

## DISCUSSION

We aimed to characterize how different alpha networks modulate perceptual decision-making. We used TDE-HMM to identify large-scale networks in a data-driven manner and focused on networks exhibiting clear oscillatory alpha activity. We linked network alpha activity to participants’ perceptual decisions using a GLMM approach and showed that ongoing alpha activity in the visual and sensorimotor networks predicts perceptual sensitivity and criterion, respectively. Namely, distinct networks exhibiting oscillatory alpha activity modulate different aspects of perceptual decision-making.

Our results not only replicate but also extend our previous ROI-based findings linking alpha activity to perception^15^. The observed sensitivity (*d’*) modulation by alpha activity in the visual network is consistent with our previous report. Here, we extend this finding by showing that this modulation was only present at times when the visual state was on, that is, when it dominated. The observed criterion (*c*) modulation by alpha activity in the sensorimotor network was unanticipated, though it is consistent with our previous observation that increased pre-stimulus alpha power in the left somatosensory and motor areas led to more liberal criterion. This modulatory effect was considered elusive in our previous report, because it was observed very briefly at ∼500 ms before stimulus onset, and only in the conservative condition.

The four alpha states identified here exhibit different spectral profiles. Notably, the peak frequency of the alpha oscillatory components was different between the visual and sensorimotor states, in line with the idea that different alpha generators contribute to these different networks. These results are consistent with findings by Rodriguez-Larios and colleagues (2022) using ICA to differentiate different alpha oscillatory components, in that the more posterior alpha network (i.e., the visual network in our case, or Alpha2 in Rodriguez-Larios et al. (2022)) showed a higher peak frequency. It is an open question whether the frequency difference between these two alpha generators is universal and generalizable.

Our current results provide important insights into how ongoing (or pre-stimulus) alpha activity modulates perception. Previous work attempting to address this question has typically used an ROI-based approach, where paired contrasts (e.g., hit vs. miss) are applied to sensor-level M/EEG data to chart the spatial extent of the modulation^11–13,31,32^. The resulting sensor-level significant clusters typically involve many sensors covering occipital, temporal and parietal areas. However, few studies have sought to distinguish whether one or more sources contribute to these dynamics, even though it has been widely accepted that more than one alpha source (or network) exists in the human brain. Therefore, the mixed findings on how alpha activity modulates perception could be due to different studies examining different alpha networks of distinct perceptual relevance. The current findings, by explicitly teasing apart different alpha networks and modeling their moment-to-moment dominance, provide concrete support for this explanation.

The idea that alpha activity enables information routing via gain control has been challenged by recent studies, in which the extents of attention-related alpha modulations were found to be independent of the sensory gain modulations of stimulus-related responses^33–35^. Therefore, instead of gain control, these researchers suggested that alpha activity acts to gate information, regulating what information passes through the processing hierarchy and what gets blocked. Our current findings, which demonstrate that distinct alpha networks modulate different aspects of perception, support both the gain control and gating hypotheses. Specifically, our observation that the visual network influences perceptual sensitivity to faint visual stimuli aligns with the gain control hypothesis. Additionally, our finding that the sensorimotor network modulates decision criterion is consistent with the gating hypothesis. In other words, alpha activity in the sensorimotor network does not correlate with the precision of stimulus encoding, but rather with how sensory information is thresholded and selected for interpretation. It is therefore conceivable that the two proposed mechanisms are not mutually exclusive but can co-exist and reflect distinct roles of different networks. Yet it should be noted that, as we did not measure sensory gain directly from the brain and correlate it with alpha activity of interest, our results do not provide direct evidence for this proposition.

The network approach is key to our current study. Eight spectrally and spatially distinct networks were identified in a data-driven way using TDE-HMM. Note that HMM assumes mutual exclusivity of states, and one may ask whether this assumption holds true for real brain activity and if not, to what extent such an assumption biases the results. In fact, the TDE-HMM approach has been validated against other methods^36^, and it was shown that similar results are obtained without the exclusivity assumption. Moreover, networks revealed by HMM in the current study share similar spatial topographies and connectivity profiles as those found in previous studies applying TDE-HMM to resting-state and task-based MEG data^20,29,36^, further reinforcing the robustness of our findings.

To summarize, we demonstrate that different alpha networks modulate different aspects of perceptual decision-making. These results provide crucial insights towards resolving previous conflicting findings on the perceptual relevance of alpha activity and spearhead an important paradigm shift—from regions to networks—in studying oscillatory dynamics in the human brain.

## RESOURCE AVAILABILITY

All data and code for stimulus presentation are available from the Donders Repository at https://doi.org/10.34973/w1k5-sm41. Analysis code to reproduce the current results can be accessed via GitHub repository https://github.com/yingjoeyzhou/analysis-predalpha-dynamics.

## ACKNOWLEDGMENTS

This study was supported by the Netherlands Organization for Scientific Research Rubicon Grant (019.222SG.003) awarded to Y. J. Zhou, and NIH grant R01-MH123679 awarded to S. Haegens. We thank Chetan Gohil for helpful discussions.

## AUTHOR CONTRIBUTIONS

Conceptualization, Y.J.Z. and S.H.; Methodology, Y.J.Z., M.W.J.v.E., and S.H.; Investigation, Y.J.Z., M.W.J.v.E., and S.H.; Writing – Original Draft, Y.J.Z.; Writing – Review & Editing, Y.J.Z., M.W.J.v.E., and S.H.; Funding Acquisition, Y.J.Z. and S.H.; Resources, Y.J.Z., M.W.J.v.E., and S.H.; Supervision, S.H.

## DECLARATION OF GENERATIVE AI AND AI-ASSISTED TECHNOLOGIES

During the preparation of this work, the author(s) used chatGPT for grammar check and text polishing. After using this tool or service, the author(s) reviewed and edited the content as needed and take(s) full responsibility for the content of the publication.

## STAR Methods

### EXPERIMENTAL MODEL AND STUDY PARTICIPANT DETAILS

This study used published data from our previous work^15^. In brief, we used MEG to record brain activity from 32 healthy participants while they performed a visual detection task. In each trial, either a grating or a noise patch was presented briefly for 16.7 ms, followed by a 100-ms high contrast mask, and the participant had to report whether they saw a grating or not (Figure 1A). We introduced criterion shifts using a “priming” method. Specifically, each block started with 32 “priming” trials, where supra-threshold gratings were presented in 20% of the trials in the conservative condition, and in 80% of the trials in the liberal condition (Figure 1B). These priming trials were followed by 80 main trials, where threshold-level gratings were presented in 50% of the trials, regardless of whether the current block was of the conservative or liberal condition. The experiment consisted of eight blocks in total. Our analyses primarily focused on the main trials, where the bottom-up sensory inputs were perfectly matched between conditions.

## METHOD DETAILS

### MEG preprocessing

MEG data were preprocessed off-line using the python-based toolboxes MNE^37^ and osl-ephys^38,39^. The raw continuous data were down sampled to 250 Hz, bandpass filtered between 0.25 and 125 Hz, and notch filtered to remove line noise (50 Hz) and its harmonic (100 Hz). Data segments and channels containing outlier values were automatically detected and removed using osl-ephys functions “detect_badsegments” and “detect_badchannels” with default settings before entering independent component analysis (ICA). ICA components were visually inspected, and components representing eye- and heart-related artifacts were projected out of the data^40^.

MRI data were co-registered to the CTF coordinate system using the fiducial coils and the digitized scalp surface. Volume conduction models were constructed based on single-shell models^41^ of individual participants’ anatomical MRIs for 29 participants, or a template MRI for three participants because their MRI were not available. Sensor data were projected onto an 8-mm grid, with dipole positions constructed using a template brain (MNI152). To ensure comparability of source reconstructions across participants, we first warped each participant’s anatomical MRI to the template brain, and then applied the inverse warp to the grid. The grid was further grouped into 52 parcels based on a refined version of the Glasser atlas^42,43^.

### Source reconstruction

We used the LCMV beamformer approach^44^ to estimate brain activity at the source level. The data covariance matrix was computed on the preprocessed continuous data. The brain activity time course of an anatomical parcel was computed by taking the first principal component of the time series over all dipole positions within the parcel. The resulting parcel time-courses were orthogonalized using Multivariate Symmetric orthogonalization^45^. Finally, the sign ambiguity resulting from the beamformer approach was resolved by applying a sign-flipping algorithm based on lagged partial correlations^20^.

### Hidden Markov modeling

We used the time delay embedded hidden Markov model (TDE-HMM)^20^ to find large-scale brain networks in a data-driven way. The HMM is a generative model, which assumes that the observed data are generated by a finite number (*K*) of recurring, transient, and mutually exclusive hidden states (Figure 1C). The data of high spatial dimension at each time point were associated with one of the states. Each state can be characterized by a spatial-spectral profile, in terms of power spectral density and within-area and between-areas connectivity profiles. The occurrence of states is assumed to be Markovian, namely, the current state only depends on the immediate previous state. The recurrence and transitions of states are captured by the transition probability. We specified 8 states and 15 embeddings (corresponding to lags of -28 to +28 ms, including lag = 0 ms), and used a multivariate Gaussian observation model with zero mean for the model fit. To ensure the stability of HMM results across different inference runs, we performed the TDE-HMM model fit for ten times and used results with the lowest free-energy for further analysis.

The primary output of this analysis is the posterior probability of each state at each time point (commonly known as Gammas), which was subsequently binarized to derive the hidden state time course (also known as the Viterbi path). Both metrics provided a dynamic latent representation of the observed data over time. Importantly, the HMM was fitted to the continuous data, without any knowledge of the task-structure. The resulting gammas were then epoched post-hoc according to the trial information (from -1 to 1 s, locked to the stimulus onset of each main trial), averaged, and baseline corrected (with a 100-ms pre-stimulus window locked to stimulus onset), to obtain the state’s stimulus-evoked responses.

We computed the spectral information (power spectral density and coherence) for each state using a multitaper approach (taper window length of 2 s, frequency range of [1, 45] Hz, and frequency resolution of 0.5 Hz). Seven Slepian tapers were fitted to the parcel time series data, conditioned on the on-off status of each state, resulting in participant-, parcel-, and state-specific PSDs and cross PSDs. We then used the PSDs to compute power maps and the cross PSDs to compute coherence networks, following the approach proposed earlier^46^. Furthermore, we applied non-negative matrix factorization (with two modes) to the stacked participant- and state-specific coherence spectra to identify common frequency bands of coherent activity. These steps were implemented using the osl-dynamics toolbox^36^.

### Spectral analysis on network activity

To identify and select alpha networks, we specifically inspected the alpha-band power and coherence maps of the resulting states. Parcels of 0.25% strongest connectivity were defined as the most critical nodes of the network, resulting in 6-7 critical nodes per network. Note we considered selecting a handful of nodes, instead of taking all parcels, a crucial step to improve the signal-to-noise ratio in estimating network-level alpha activity (i.e., *P*_*t*,*k*_ in the GLMM). This is because it prevents the variability between areas—caused by averaging across highly heterogeneous parcels—from obscuring the trial-by-trial activity fluctuations. The network-level PSDs were defined as the averaged PSD across the corresponding critical nodes, computed using parcel time series data at time points when the state was on.

We defined the network-level alpha power time course as the averaged alpha power time courses across the selected critical nodes (Figure 1D). The alpha frequency of interest was defined for each participant and for each network by applying the FOOOF algorithm^47^ to the corresponding PSDs, to account for inter- and intra-subject variability^48^. Settings for the algorithm were as follows: peak width limits: [0.2, 12]; max number of peaks: 3; minimum peak height: 0.3; peak threshold: 2; and aperiodic mode: ‘fixed’. Power spectra were parameterized across the frequency range of 1 to 45 Hz. Based on the algorithm’s output (i.e., the fitted aperiodic and periodic/oscillatory components), we defined the individual alpha peak as the oscillatory component’s extracted center frequency that falls within the 7-14 Hz band, or as the corresponding group average when no clear oscillatory component was identified within the above-mentioned band. In rare cases where more than one extracted peak frequency fell within the 7-14 Hz band, the lower frequency was used. With these alpha peak frequencies at hand, the alpha power time courses were computed by first applying the first taper of the Slepian sequence and then a fast Fourier transform to short sliding windows of the critical nodes’ time series data (500 ms in length, sliding in 40 ms steps, ±3 Hz spectral smoothing).

### Generalized linear mixed models (GLMM)

We built generalized linear mixed models to link alpha activity to participant’s behavioral responses. We focused on sensitivity (*d’*) and criterion (*c*), key measures defined by Signal Detection Theory (SDT). Assuming equal variance for internal signal and noise distributions, we have:

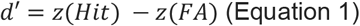

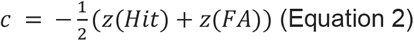

where *z*(*Hit*) and *z*(*FA*) denote the inverse of the standard normal cumulative distribution function evaluated at the given hit and false alarm (FA) rate. Higher *d’* suggests higher sensitivity, and higher *c* suggests more conservative in reporting “target present”.

To quantify the relationship between neural activity and behavioral outcomes, we adapted and extended the GLM formulation of signal detection theory^49^. Concretely, participant’s perceptual report in each trial is determined by both the stimulus and neural factors:

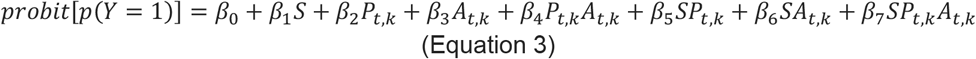

where *Y* denotes the participant’s perceptual report (“present” vs. “absent”, 1 for “present” and 0 for “absent”), *S* denotes the grating presence (present vs. absent, 1 for present and 0 for absent), *A*_*t*,*k*_ denotes the on-off status of network *k* at time *t*, and *P*_*t*,*k*_ denotes the alpha power of network *k* estimated at time *t*, computed by averaging the alpha power estimates at time *t* across the critical nodes/parcels of network *k* (Figure 1D). We used a sliding window of 500 ms to estimate alpha power, and aligned our definition of *A*_*t*,*k*_ for simplicity. Namely, if the *k*th state is on at any time point within the 500-ms sliding window centered at time *t*, then *A*_*t*,*k*_ = 1, otherwise *A*_*t*,*k*_ = 0. According to SDT, we have:

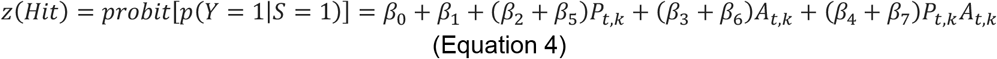

and

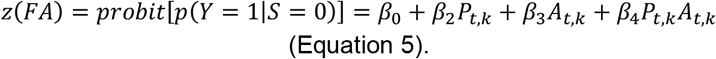

Replacing *z*(*Hit*) and *z*(*FA*) in Equation 1 with Equations 4 and 5 results in:

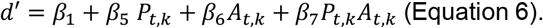

Letting *A*_*t*,*k*_ = 1 and *A*_*t,k*_ = 0 in the above equation result in:

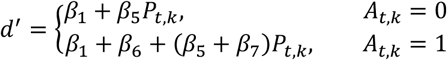

where the coefficients *β*_*d*′,*A*=1_ = *β*_5_ + *β*_7_ and *β*_*d*′,*A*=0_= *β*_5_ quantify how power changes modulate perceptual sensitivity when the state is on and off, respectively. Furthermore, the coefficient of the interaction term (i.e., *β*_7_) denotes whether power changes modulate perceptual sensitivity differently when the state is on vs. when it is off.

Similarly, replacing *z*(*Hit*) and *z*(*FA*) in Equation 2 with Equations 4 and 5 results in:

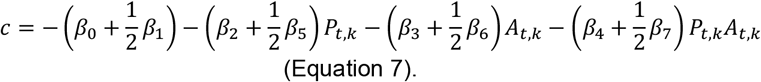

Letting *A*_*t*,*k*_ = 1 and *A*_*t*,*k*_ = 0in the above equation result in:

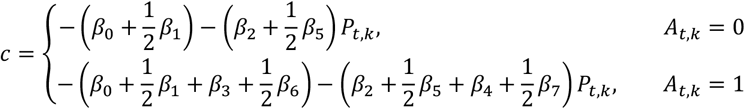

The coefficients 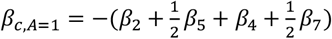 and 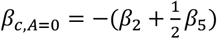 quantify how power changes modulate perceptual criterion when the state is on and off, respectively. And the coefficient of the interaction term (i.e., 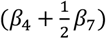 denotes whether power changes modulate perceptual criterion differently when the state is on vs. when it is off. We fit the following mixed-effects model to the data (in Wilkinson notation): *Y ∼ 1+S*P*A + (1+S* | *subject:condition)*, where *Y, S, P*, and *A* correspond to the *Y, S, P*_*t*,*k*_, and *A*_*t*,*k*_ terms of Equation 3, and *subject* and *condition* are categorical variables denoting the participant and condition that a particular trial belongs to. By modelling the intercept and slope of *S* as random effects, we accounted for differences between individuals and between conditions in their condition-specific criterion and sensitivity levels (especially given that the conservative and liberal conditions were associated with significantly different criteria). We z-scored the log-transformed trial-by-trial alpha power of interest before feeding it to the model, to reduce between-subjects heterogeneity in alpha power estimates and to enhance interpretability of the resulting model. This analysis was implemented using MATLAB’s “fitglme” function.

## QUANTIFICATION AND STATISTICAL ANALYSIS

We used cluster-based permutation tests^50^ to establish statistical significance. For time series data, clustering took place for neighboring time points where the *F* values (or *T* values) corresponded to *p* values smaller than 0.05 (uncorrected). The *F* values (or *T* values) at different time points within the cluster were summed and later used as the cluster-level test statistics. Permutation was performed by shuffling the labels of the real data and recalculating the cluster-level test statistics, to obtain a reference distribution of cluster-level maximum test statistics. Finally, the cluster-level test statistics of the real data were evaluated against the reference distribution, to obtain the statistical significance of each cluster. For the stimulus-evoked responses resulting from the HMM analysis, two-tailed paired t-tests (against zero) were used to obtain *T* values univariately at each time point. Permutation was obtained by randomly shuffling the condition labels of the stimulus-evoked responses 5000 times, or by shuffling the real data with time series of zeros 1000 times, to establish statistical significance of stimulus-evoked response (stimulus vs. baseline). For the beta coefficient time courses resulting from the GLMM analysis, the corresponding *F* values given by the GLMM were used for clustering. Permutation was obtained by refitting the GLMMs using shuffled data, in which the alpha power time courses across trials were shuffled 400 times in a within-subject manner.

