## Supplementary material for "Distinct Alpha Networks Modulate Different Aspects of Perceptual Decision-Making": Figure S1, Figure S2

**SUPPLEMENTARY FIGURES**

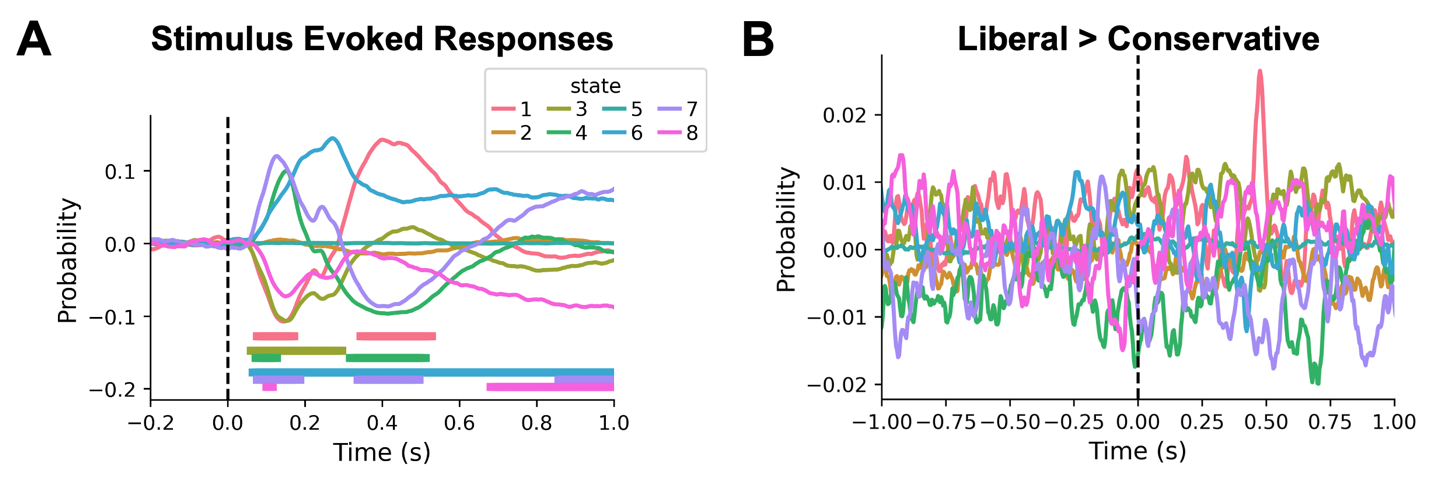

**Figure S1. (A)** State time courses epoched around the presentation of stimulus (onset denoted by the dashed black line). Horizontal bars indicate time points that contributed to the significant clusters. **(B)** Difference state time courses between the liberal and conservative conditions.

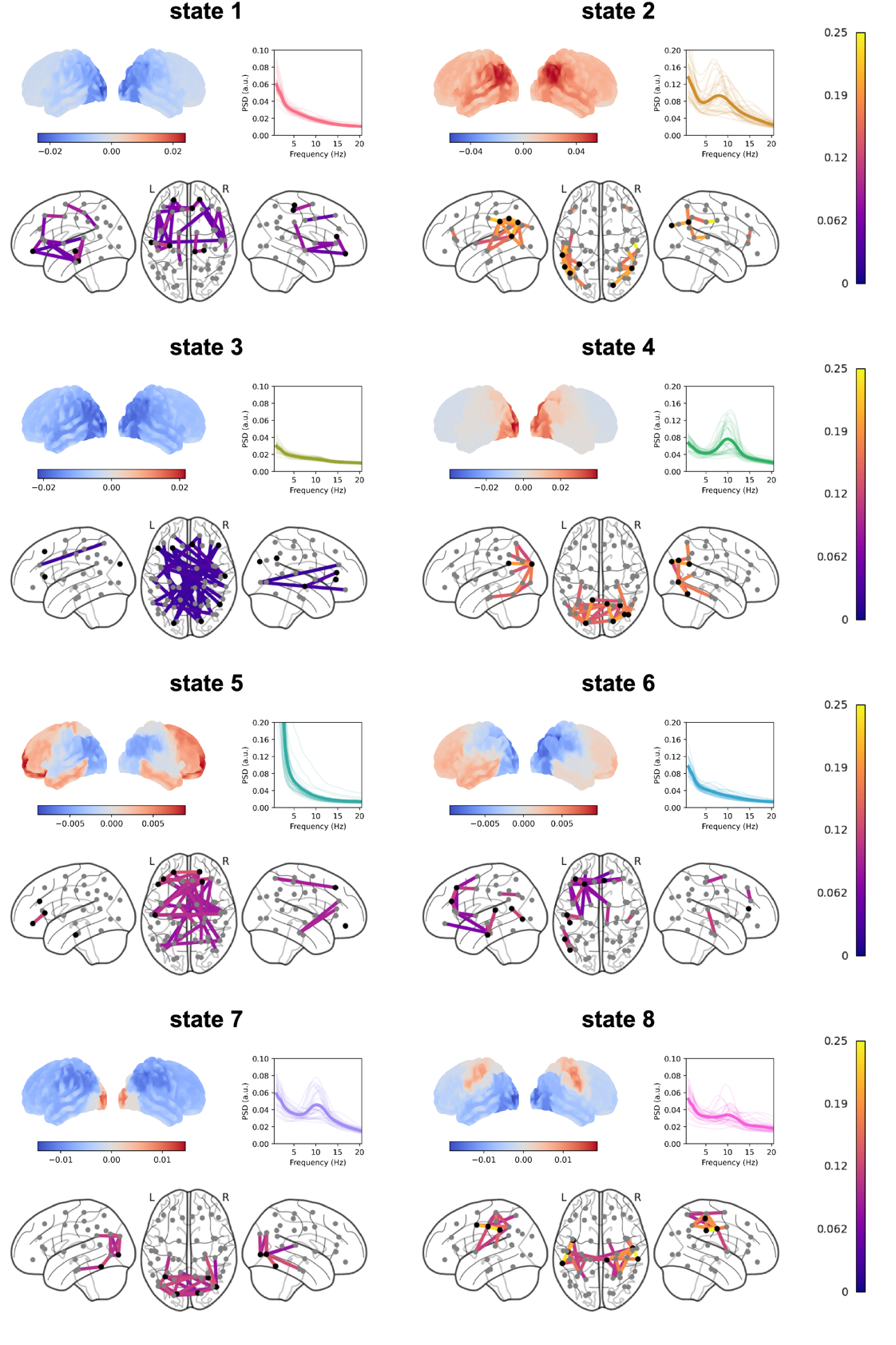
**Figure S2.** For each state, the spatial distribution of power and coherence were estimated for frequencies in the alpha range. Sub-panels are organized similarly to Figure 2; the PSD graph (top right) shows the state-specific PSD (and the corresponding standard error, shaded area) estimated from the critical nodes (highlighted in black in the coherence network).
